# Air temperature in fringe habitats: performance of climate reanalysis on Atlantic Patagonian rocky shores

**DOI:** 10.64898/2026.02.27.708471

**Authors:** M. R. Robert, N. Pessacg, J. P. Livore, M. M. Mendez

## Abstract

Climate change and particularly the frequency and intensity of extreme events is affecting the distribution and abundance of species, with drastic consequences on ecological processes and community structure. Long-term records of environmental parameters are indispensable in climatological studies in order to better understand the processes involved. However, such data is usually unavailable for many geographic areas and certain environments, like Patagonian intertidal shores in the Southwestern Atlantic. The use of reanalysis products can help elucidate the climate of the past when *in situ* information is missing. In this work, we test the performance of reanalysis datasets in reproducing air temperature patterns and extreme hot events (heatwaves) on rocky intertidal environments of Atlantic Patagonia. Thus, we evaluate the degree of correlation between different reanalysis products and air temperature data from loggers placed on rocky shores. We also test whether those products accurately detect the duration, frequency and number of heatwaves and look for historical trends in their features. Our results showed that reanalysis products perform well for assessing broad-scale changes in air temperature patterns. Products were also capable of detecting heatwaves, with little variation in their features for the period 1960–2024. Additionally, real-time field temperatures to which intertidal organisms are exposed were obtained for the first time in the area; reporting heatwaves events. Thereby, reanalysis products complement local data, providing key information to understand the role that temperature increases and extreme heat can have in events like mussels mass mortalities reported locally. In this sense, our results suggest that heatwaves alone wouldn’t be explaining the observed mussel losses. This work provides empiric evidence on the usefulness of reanalysis products of intertidal habitats and encourages similar approaches in order to properly understand climatological patterns that can drive ecological processes on coastal habitats.

## Introduction

In natural environments, some species are undoubtedly more significant in determining overall community structure than others. Ecologists describe these foundation species as those that have a large effect on community structure by modifying environmental conditions, species interactions, and resource availability through their presence (i.e. foundation species, following Bruno and Bertness, 2001). Mussels are a distinctive foundation species dominating temperate rocky shores worldwide (Paine and Levin, 1981; Thiel and Ullrich 2000; Bertness et al., 2006). They tend to form dense layers with practically all mid intertidal organisms unable to survive outside of the mussel bed (Bertness et al., 2006; Mendez et al., 2021). The effect of extreme hot events on these intertidal foundation species is relevant given their ecological role and their sessile nature that makes them particularly vulnerable to heat stress and desiccation (Wernberg et al., 2024). Several studies have demonstrated that current climate change has caused substantial declines in foundation species and their associated communities (reviewed by Wernberg et al. 2024). In fact, extreme temperature events have been responsible for mass mortalities in a number of foundation species on rocky shores (Garrabou et al., 2009; Harley, 2008; Hesketh and Harley, 2023; Thomsen et al. 2019).

The effects of anthropogenic actions on climate are having severe ecological and socio-economic consequences; affecting the abundance, physiology, reproduction, and distribution of species worldwide (Harley et al., 2006; Hawkins et al., 2008; Helmuth et al., 2006; Bates et al., 2018; Wernberg et al., 2024). Climate change modifies in the last decades not only average conditions but also the frequency and intensity of extreme events, which are predicted to continue to rise towards the future. In this sense, heatwaves constitute an undeniable risk to thermally sensitive marine species. Although various definitions exist, heatwaves are generally understood as periods of at least three consecutive days during which temperatures exceed a high percentile (typically the 90^th^) of the long-term climate records. These extreme events have an important role in structuring biological communities since, unlike gradual increases, the acuteness of these events may not allow organisms to acclimate (Wernberg et al., 2013; Bennet et al., 2015; Stillman, 2019; Wernberg et al., 2024). Hence, analyzing heatwave variability and trends in order to understand how potential environmental conditions will affect species is essential given the rapid and unprecedented impacts already recorded on biological communities.

Continuous and extensive records of environmental parameters are indispensable in climatological and ecological studies, as they allow the detection of climatic variations at different temporal and spatial scales (Thompson et. al, 2002; Hawkins et al., 2008; Mieszkowska et al., 2021; Bhattacharyya et al., 2025). However, this inevitably depends on the accessibility of long-term *in situ* data of those environmental parameters (Mieszkowska et al., 2021; Seidnitzer et al., 2024; Bhattacharyya et al., 2025). The availability of such information is usually restricted by different factors, principally historical and current funding and logistics. The use of reanalysis (i.e. retrospective analyses) of environmental parameter databases emerges as an alternative to this problem. Main atmospheric variables, including temperature, precipitation, and relative humidity, can be obtained for retrospective periods at high spatial and temporal resolution for the entire globe or a particular site. Briefly, climate reanalysis consists in combining precise observational data with weather prediction models (Thorne and Vose, 2010; Fujiwara et al., 2017). In the last decades, a number of climate reanalysis products have been developed incorporating subsequent improvements with some of them reaching third and fourth-generation versions (Hersbach et al., 2020; Yang et al., 2022). Previous works have shown that reanalyses performed better in temperate than in tropical and subtropical regions and they can reasonably reproduce climatological variations and trends in atmospheric variables globally (Donat et al., 2014).

Atlantic Patagonian shores are exposed to a temperate climate that includes strong dry and persistent winds, low humidity and rainfall and a large thermal amplitude. A dense matrix of the scorched mussel *Perumytilus purpuratus* dominates the physiognomy of these rocky shores (Bertness et al., 2006; Silliman et al., 2011; Livore et al., 2021). Historically, beds from this region show simple structure, uniform appearance and disturbance-generated bare space throughout the bed is strikingly rare (Bertness et al., 2006; Adami et al., 2018). However, drastic losses in cover of scorched mussels were registered at different sites after the 2019 austral summer; where mussel cover dropped from 90% to almost no living organisms (Mendez et al. 2021). This loss of scorched mussels coincided with high atmospheric temperature occurring during low tides that exposed these organisms to prolonged thermal and desiccation stress over several consecutive days.

In order to determine the relative importance of temperature increases and extreme heat in events like the loss of mussels, information on the thermal patterns to which the species has been exposed in the last decades is needed. In this context, it is important to note that in Atlantic Patagonia, Southwestern Atlantic, meteorological information is extremely scarce, with only 18% of the existing meteorological stations belonging to a national network (Almonacid et al. 2022) and virtually no coastal information. Thus, reanalysis datasets become an alternative to direct observations and could fill these information gaps, playing a crucial role in data-sparse regions with limited observation over time and space. However, they have inherent biases, making it necessary to identify the most suitable reanalysis dataset for specific areas. While the assessment of reanalysis datasets is common, how well recent generation reanalyses detect temperature trends and extreme events remains an open question. Even though reanalyses have not been designed to provide evidence at the local level, given the scarcity of data in the region and the improvement in the spatial resolution of these products, they can provide valuable information. Therefore, this study aimed to test the performance of reanalysis datasets in reproducing air temperature patterns and extreme hot events on rocky intertidal environments of Atlantic Patagonia. To do this, we first evaluate the degree of correlation between different reanalysis products and air temperature data from loggers placed on rocky shores. We then test whether the reanalysis products accurately detect the duration, frequency and number of extreme temperature events (heatwaves) and look for historical trends in heatwave frequency.

## Materials and methods

### Study sites

The analysis was conducted at two intertidal sites in northern Patagonia, Southwestern Atlantic: Punta Buenos Aires (PBA. Fig. 1) and Punta Loma (PLM. Fig. 1). At each site, temperature loggers (EnvLogger version T2.4, ElectricBlue CRL) were deployed at the mid-intertidal level. At PBA, two austral summers (2022–2023 and 2023–2024) were covered, whereas at PLM data covered three summers (2022–2023, 2023–2024, and 2024–2025). The loggers recorded temperature at hourly intervals.

**Fig. 1.**
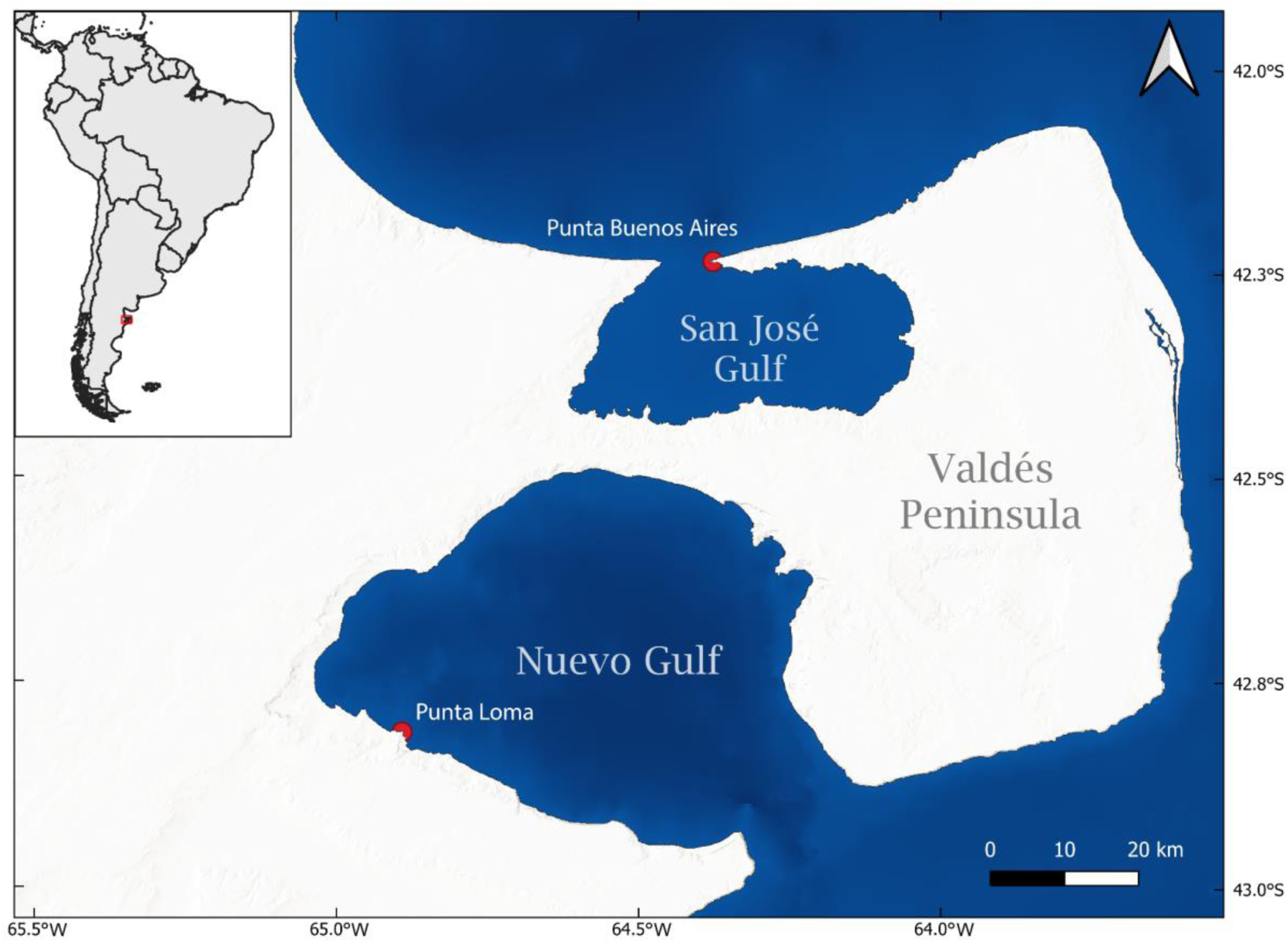
Study area and location of the intertidal sampling sites (Punta Buenos Aires and Punta Loma) in northern Patagonia, Southwestern Atlantic.

### Validation of reanalysis products against in situ intertidal temperature records

Temperature records were filtered to retain only those corresponding to periods of emersion, that is, when the sensors were exposed to air due to tidal retreat. This filtering was performed independently for each site by computing hourly tidal heights using the global harmonic model TPXO9-v5 (Egbert & Erofeeva, 2002) through the Python library **pyTMD**, which implements harmonic interpolation of tidal constituents. The four main constituents (M2, S2, K1, and O1) were extracted at each sités geographic coordinates. The resulting tidal series were validated against predictions from the Argentine Hydrographic Service (SHN) for nearby maritime ports. Based on tidal height and the vertical position of the loggers relative to mean sea level, each record was classified as either emersion or immersion.

Hourly series of 2-m air temperature (t2m), representing near-surface atmospheric conditions, and skin temperature (skt), corresponding to the radiative temperature of the surface, were extracted from the ERA5 reanalysis (Hersbach et al., 2022), which provides global data at ∼0.25° spatial resolution (∼25km), and from ERA5-Land (Muñoz-Sabater et al., 2019; 2021), which offers land-surface variables at a higher spatial resolution of ∼0.1° km (∼10km). Both datasets are produced by the European Centre for Medium-Range Weather Forecasts and distributed through the Copernicus Climate Change Service Climate Data Store. Additionally, data were obtained from the MERRA-2 reanalysis (Modern-Era Retrospective Analysis for Research and Applications, Version 2; Gelaro et al., 2017), which has a spatial resolution of approximately 0.5° (∼50 km), for the same sites and periods as the logger deployments. For each site, the nearest available grid cell in each reanalysis dataset was selected. The commonly used average over the four adjacent grid cells was not applied, given the highly heterogeneous nature of the intertidal environment. Including adjacent cells would have added cells that correspond to sea water which would increase variability and introduce unwanted errors. All reanalysis series were subsequently filtered to retain only the records corresponding to time of emersion.

Because reanalysis datasets exhibit systematic errors (Hurtado et al., 2024), a bias correction was applied using the Quantile Mapping (QM) method implemented in the *xsdba* Python package. This non-parametric approach adjusts the distribution of modeled values to match the empirical distribution of observed temperatures. As a reference, only the *in situ* logger temperatures recorded during emersion were used, since they represent the actual thermal conditions experienced by intertidal organisms. Although the observational record covers only two or three summer seasons, applying QM remains justified because it provides the only available basis for aligning the distributional characteristics of the reanalysis products with the conditions actually measured in the field. In this context, the aim of QM is not to reconstruct a full climatology but to correct systematic biases within the period for which observations exist, thereby improving the physical consistency between modeled and observed temperatures. For this reason,t he correction was implemented independently for each site and summer season, mapping the empirical percentiles of each reanalysis product (ERA5, ERA5-Land, and MERRA-2) to the corresponding percentiles of the observed distribution. This approach corrects systematic biases across the entire distribution, including extreme values relevant for heatwave detection.

For each site and summer season, agreement metrics were computed between logger and reanalysis data: mean bias (Bias), mean absolute error (MAE), and root mean square error (RMSE). The performance of corrected and raw datasets was then evaluated using violin plots to compare error distributions, and by computing Pearson’s correlation coefficients between observed and reanalysis temperatures.

To evaluate systematic biases and distributional discrepancies between *in situ* intertidal temperatures and reanalysis products, we constructed Quantile–Quantile (Q–Q) plots for each site and summer season. Hourly emersion temperatures recorded by the loggers were treated as the observational reference. For the three reanalysis dataset evaluated, we compared both the original (raw) temperatures and the corrected series obtained through QM. The Q–Q procedure consisted of computing quantiles of the reanalysis data and plotting them against the corresponding quantiles of the logger temperature distribution.

### Heatwave event detection capacity of reanalyses

Heatwaves were defined as events of at least three consecutive days in which at least one hourly temperature measurement recorded during emersion per day exceeded the 90th percentile of logger temperatures at each site. Detection was implemented using a Python script that identified calendar-day sequences containing at least one hourly value above the mentioned thermal threshold. Heatwaves detected from logger data were treated as the observational reference against which ERA5, ERA5-Land, and MERRA-2 detections (both raw and QM-corrected) were compared using the same criteria. This approach allowed assessing the detection skill of each reanalysis product, with and without bias correction.

### Historical trends in heatwave frequency

Using the dataset generated in the previous section, historical trends in maximum temperatures and heatwave occurrence in the intertidal zone of the Southwestern Atlantic were evaluated. Daily maximum temperatures during emersion for the period 1960–2024 were analyzed for the PLM site, following the same emersion-filtering methodology described above. No quantile mapping correction was applied to the long-term series because the available in situ observational period was insufficient to robustly calibrate the model’s distribution (Maraun, 2013).

Long-term monotonic trends in several heatwave metrics were assessed: the annual number of events, the mean and total event duration (days), the mean maximum temperature, and the mean event temperature. The mean maximum temperature was calculated as the annual average of the maximum temperature reached during each individual heatwave event. The Mann–Kendall test—widely used in climatology to detect monotonic trends—was applied, and Sen’s slope was calculated to quantify the magnitude of the changes and express them as decadal trends. For comparative purposes, differences between two periods (1961–1990 vs. 1991–2020) were evaluated to estimate relative changes in heatwave metrics. All analyses were performed using **scipy.stats** library in Python.

Additionally, to detect possible abrupt changes (breakpoints) in the maximum temperature series, the non-parametric Pettitt test (Pettitt, 1979) was applied. This test estimates the U statistic and an associated p-value, identifying the year of a potential shift and the mean values before and after the change. The **pyhomogeneity** library (Hussain et al., 2023) was used for this analysis. To assess thermal trends within the periods defined by the detected breakpoint, we applied the non-parametric Mann–Kendall trend test to the series of annual maximum temperatures. Analyses were conducted independently for the periods before and after the breakpoint in order to evaluate potential differences in trend behavior across the two regimes.

## Results

### Validation of reanalysis products against in situ intertidal temperature records

The comparison between hourly temperature records obtained from in situ loggers and reanalysis products, based on correlation analysis, mean absolute error, root mean square error, and bias metrics, showed an overall adequate performance at both sites (Table 1).

**Table 1.** Performance metrics of reanalysis datasets evaluated against in situ logger temperature records at two intertidal sites in northern Patagonia: Punta Buenos Aires (PBA) and Punta Loma (PLM). Bias, mean absolute error (MAE), and root mean square error (RMSE) are expressed in °C, and Pearson’s correlation coefficient (Corr.) is reported as a measure of temporal agreement. *N* indicates the number of hourly observations. Dataset names including the suffix “_qm” correspond to time series corrected using empirical quantile mapping. Bold values indicate the highest correlation coefficients and the lowest MAE and RMSE for each site.

| Site | Variable | Corr. | MAE | RMSE | Bias | N |
| --- | --- | --- | --- | --- | --- | --- |
| PBA | t2m_ERA5 | 0.84 | 2.68 | 3.57 | 0.18 | 7058 |
| PBA | t2m_ERA5_land | 0.84 | 2.78 | 3.64 | 0.37 | 7058 |
| PBA | t2m_MERRA2 | 0.77 | 3.15 | 4.20 | -0.05 | 7058 |
| PBA | skt_ERA5 | **0.91** | 2.24 | 2.93 | 0.51 | 7058 |
| PBA | skt_ERA5_land | **0.91** | 3.05 | 4.48 | 2.03 | 7058 |
| PBA | skt_MERRA2 | 0.74 | 3.50 | 4.67 | 0.37 | 7058 |
| PBA | t2m_ERA5_qm | 0.83 | 2.59 | 3.74 | 0.00 | 7058 |
| PBA | t2m_ERA5_land_qm | 0.83 | 2.64 | 3.75 | 0.00 | 7058 |
| PBA | t2m_MERRA2_qm | 0.76 | 3.23 | 4.52 | 0.00 | 7058 |
| PBA | skt_ERA5_qm | **0.91** | **1.83** | **2.76** | 0.00 | 7058 |
| PBA | skt_ERA5_land_qm | **0.91** | 1.92 | 2.80 | 0.00 | 7058 |
| PBA | skt_MERRA2_qm | 0.76 | 3.22 | 4.45 | 0.00 | 7058 |
| PLM | t2m_ERA5 | 0.88 | 3.00 | 3.87 | 1.31 | 6283 |
| PLM | t2m_ERA5_land | 0.87 | 3.06 | 3.90 | 1.22 | 6283 |
| PLM | t2m_MERRA2 | 0.91 | 2.44 | 3.11 | 0.53 | 6283 |
| PLM | skt_ERA5 | **0.95** | 3.82 | 5.09 | 2.58 | 6283 |
| PLM | skt_ERA5_land | 0.92 | 4.09 | 5.49 | 2.85 | 6283 |
| PLM | skt_MERRA2 | **0.95** | 2.12 | 2.78 | 1.23 | 6283 |
| PLM | t2m_ERA5_qm | 0.88 | 2.68 | 3.51 | 0.00 | 6283 |
| PLM | t2m_ERA5_land_qm | 0.87 | 2.83 | 3.67 | 0.01 | 6283 |
| PLM | t2m_MERRA2_qm | 0.91 | 2.33 | 3.05 | 0.01 | 6283 |
| PLM | skt_ERA5_qm | **0.95** | 1.81 | 2.40 | 0.01 | 6283 |
| PLM | skt_ERA5_land_qm | 0.92 | 2.21 | 2.85 | 0.01 | 6283 |
| PLM | skt_MERRA2_qm | **0.95** | **1.73** | **2.28** | 0.01 | 6283 |

The correlation analysis between reanalysis products and logger temperature data revealed statistically significant relationships across all models and sites (p < 0.0001), with Pearson correlation coefficients indicating a strong agreement between reanalysis datasets and field measurements (Table 1). At PBA, the skt showed higher correlation coefficients compared to the t2m. ERA5 and ERA5-Land performed consistently for both raw and QM-corrected data. Specifically, the raw ERA5 skin temperature achieved the highest correlation, while MERRA-2 showed comparatively lower but still strong correlations. The QM correction produced only minimal changes in the Pearson r values, typically smaller than 0.02 compared to the raw data. At PLM, distinct patterns emerged: MERRA-2 performed better across both temperature variables. Across both sites, skt consistently outperformed the t2m.

At PBA, the smallest errors were observed for the QM ERA5 skt, which also exhibited a high correlation with the *in situ* measurements. MERRA-2 skt product exhibited a reduced bias, but relatively high absolute and quadratic errors. At PLM, the best agreement was obtained with MERRA-2 skt, which showed the lowest error values and a moderate positive bias. Overall, ERA5 skt and MERRA-2 skt emerged as the best-performing products for both the raw reanalysis data and the datasets corrected using quantile mapping.

The violin plots showed distributions of individual errors between temperatures recorded by loggers and those estimated by the different reanalysis products (Fig. 2). At PBA, the raw products showed wide distributions, with errors reaching up to ±10 °C. The ERA5 skt showed a comparatively narrower spread, whereas ERA5-Land skt displayed longer tails toward positive errors, indicating a tendency to overestimate temperatures. In contrast, MERRA-2 skt showed extended tails toward negative errors, reflecting a systematic underestimation. At PLM, the raw products also presented asymmetric error distributions, with pronounced positive biases in the ERA5 and ERA5-Land skt. Conversely, MERRA-2 skt exhibited the most concentrated distribution and the smallest range of variation, consistent with the lower MAE and RMSE values reported in Table 1.

**Fig. 2.**
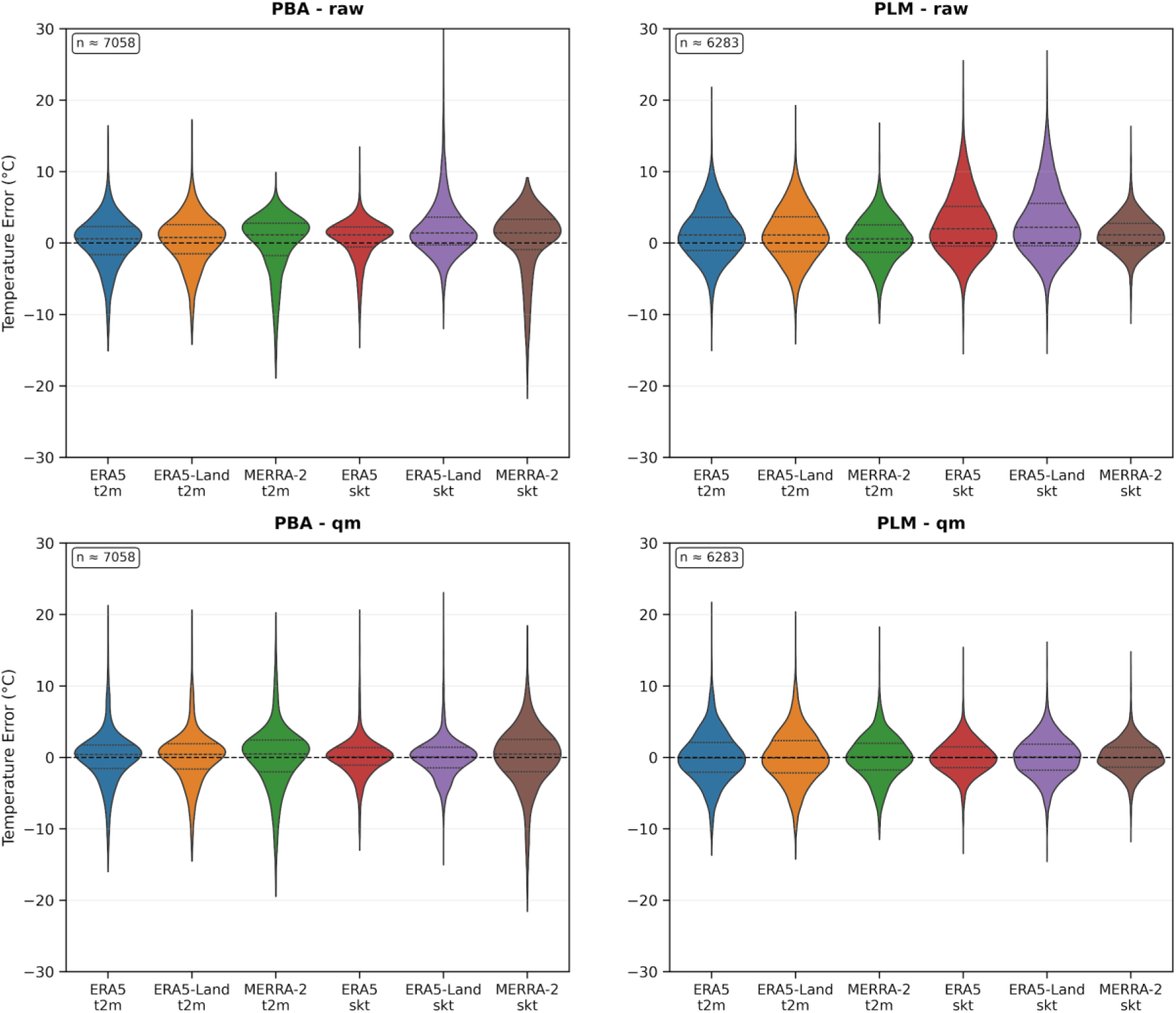
Violin plots of hourly temperature errors (reanalysis minus in situ logger) at two intertidal sites in northern Patagonia: PBA (left) and PLM (right). Panels show raw reanalysis data (top) and values after Quantile Mapping (QM) correction (bottom). Horizontal dashed lines indicate zero error, while solid lines within violins denote the 25th, 50th, and 75th percentiles. The number of observations (n) used in each comparison is shown in the upper-left corner of each panel.

Applying the Quantile Mapping (QM) correction resulted in a clear reduction in both error spread and bias at both sites (Table 1 and Fig. 2). Overall, beyond differences in mean errors, there are notable contrasts in the shape and dispersion of the distributions. The QM correction improved the representation of variability and reduced the magnitude of extreme errors.

The Q–Q plots revealed differences in how each reanalysis product captured the distribution of intertidal temperatures (Fig. 3 and Fig 4). Across both sites, the raw datasets showed substantial departures from the 1:1 line, particularly in the upper quantiles, where reanalyses tended to underestimate or overestimate the highest emersion temperatures.

**Fig. 3.**
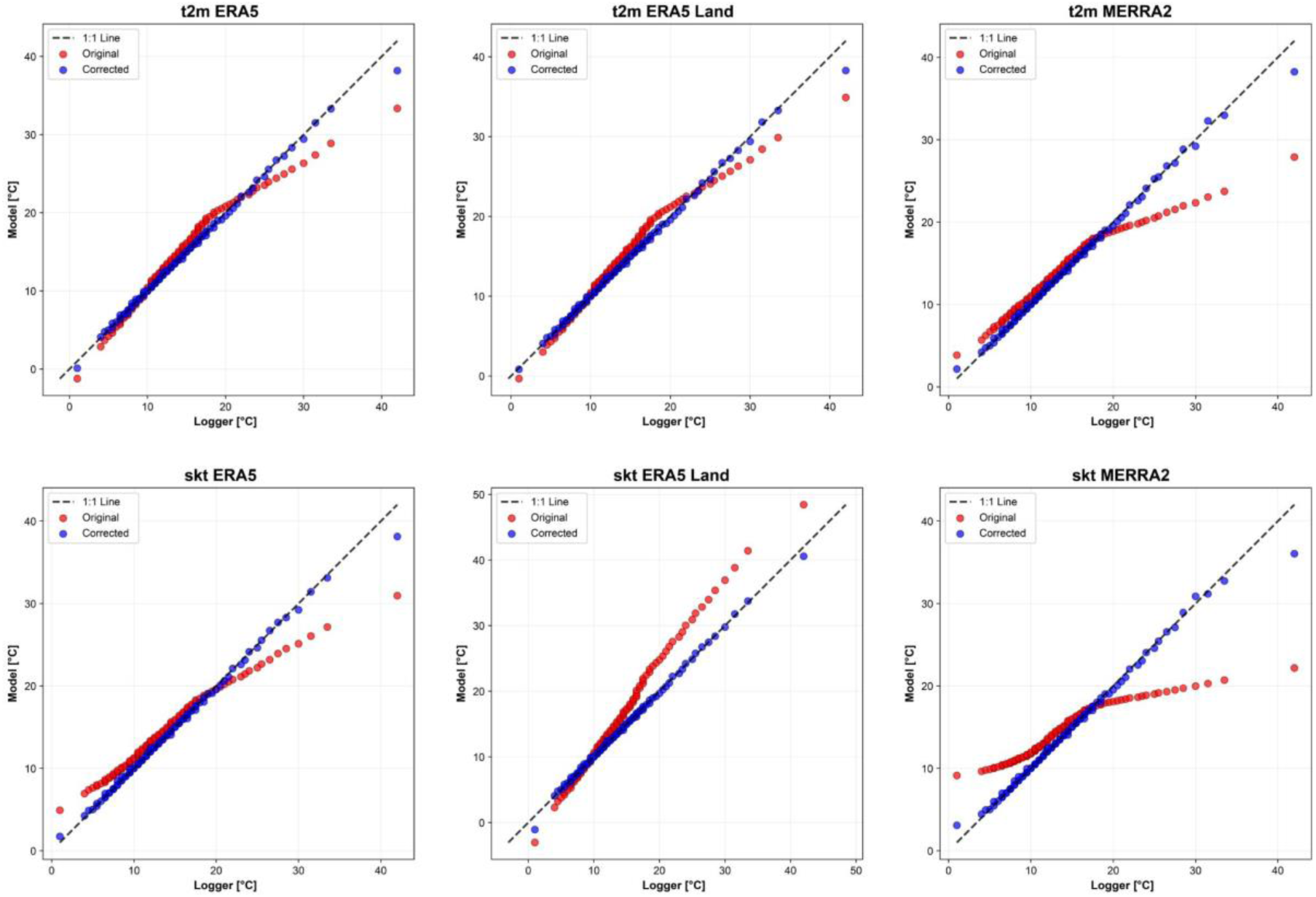
Q–Q plots comparing modeled temperatures from three reanalysis products (ERA5, ERA5-Land, and MERRA-2) with *in situ* logger observations at the PBA site. For each dataset, the original (red) and Quantile-Mapping–corrected (blue) temperatures are shown against the 1:1 line (dashed).

**Fig. 4.**
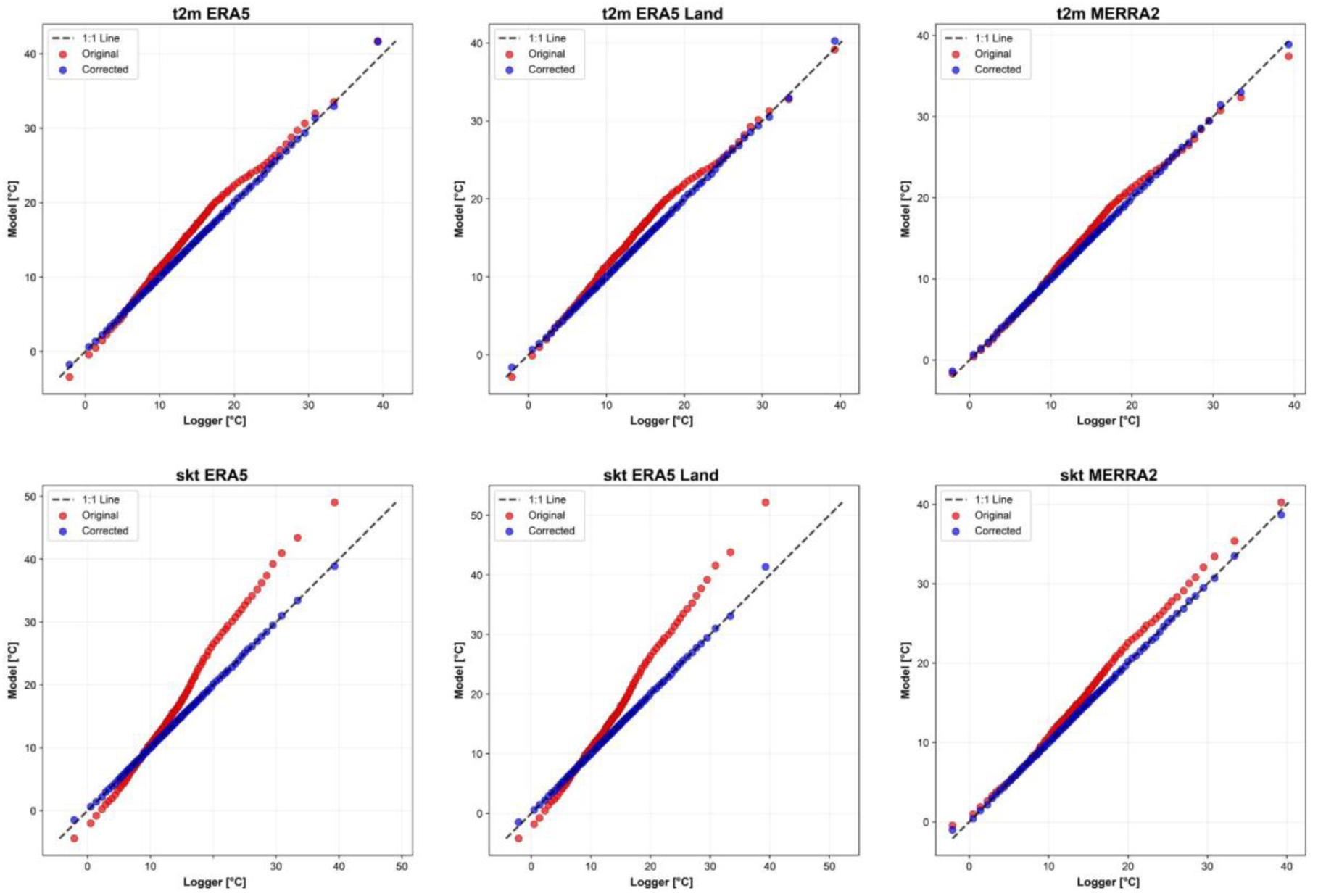
Q–Q plots comparing modeled temperatures from three reanalysis products (ERA5, ERA5-Land, and MERRA-2) with *in situ* logger observations at the PLM site. For each dataset, the original (red) and Quantile-Mapping–corrected (blue) temperatures are shown against the 1:1 line (dashed).

However, in the raw series, the t2m generally showed a closer agreement with observations than the skt. In this regard, ERA5 and ERA5-Land t2m exhibited a better overall alignment across most quantiles, whereas skt—especially MERRA-2—displayed larger deviations, mainly toward the distribution tails. After applying QM, all corrected series showed a marked improvement, with points falling much closer to the 1:1 line across the full range of temperatures. High-quantile behavior was notably improved, indicating that QM successfully removed systematic distributional biases and enhanced the representation of extreme thermal conditions.

## Heatwave event detection capacity of reanalyses

The 90th-percentile threshold for logger-recorded temperatures during emersion, used to define heatwave events, was 28 °C at both PBA and PLM. Based on this threshold, six heatwaves were detected at PBA during the 2022–2023 summer (mean duration = 6.75 ± 3.18 days), and seven during the 2023–2024 summer (6.00 ± 2.94 days). At PLM, four heatwaves occurred in the 2022–2023 summer (7.00 ± 2.94 days), four in 2023–2024 (6.50 ± 3.11 days), and one in the 2024–2025 summer (6 days).

At PBA, the raw reanalysis products showed highly variable performance in heatwave detection (Fig. 5A). The t2m (ERA5 and ERA5-Land) detected only 3 of the 13 observed events (23.1% detection rate), whereas ERA5 skt detected only one event (7.7%). Conversely, ERA5-Land skt achieved a 100% detection rate of heatwaves recorded by the loggers (Fig. 5A), but overestimated the total number of events by identifying five false positives not observed in the field (Fig. 6). In accordance with this observation, ERA5-Land skt exhibited a propensity to overestimate the duration and intensity of heatwaves (Fig. 5B, C), indicating that temperatures frequently exceeded the detection threshold. This over-detection indicates that the high matching rate does not necessarily reflect an accurate representation of heatwave occurrence. Following the implementation of QM correction, there was a substantial enhancement in the performance of all products, with detection rates ranging from 69.2% to 92.3%. It is important to note that the QM corrected products demonstrated a reduction in the number of false positives, indicating a more realistic representation of extreme events. For instance, ERA5 skt QM showed a high detection rate (92.3%) with a single false positive, while ERA5-Land skt QM achieved 84.6% with one false positive.

**Figure 5.**
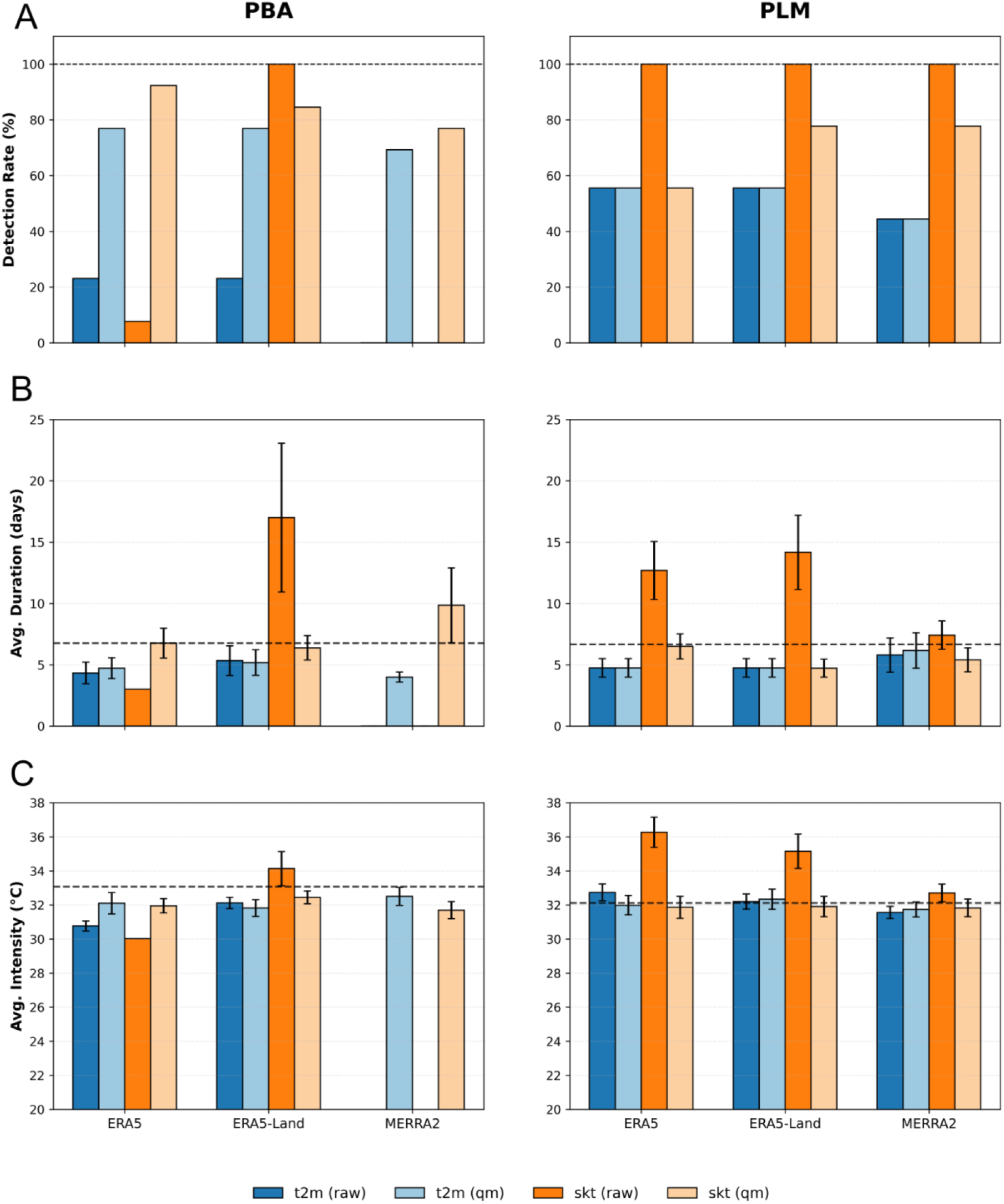
Heatwave detection metrics for PBA (left column) and PLM (right column). Blue and orange bars correspond to t2m and skt, respectively; darker and lighter shades indicate raw and quantile-mapped (QM) data. (A) Detection Rate (%), defined as the percentage of observed heatwaves correctly identified by each reanalysis product using a 2-day overlap window. (B) Mean heatwave duration (days), where the dashed horizontal line indicates the average duration observed by the *in situ* logger. (C) Mean heatwave intensity (°C), calculated as the average of daily maximum temperatures over the duration of each event; the dashed line represents the corresponding logger-based reference. Error bars in (B) and (C) indicate the standard error of the mean. All metrics were computed using a strict temperature threshold of 28.0 °C sustained for a minimum of three consecutive days.

**Fig. 6.**
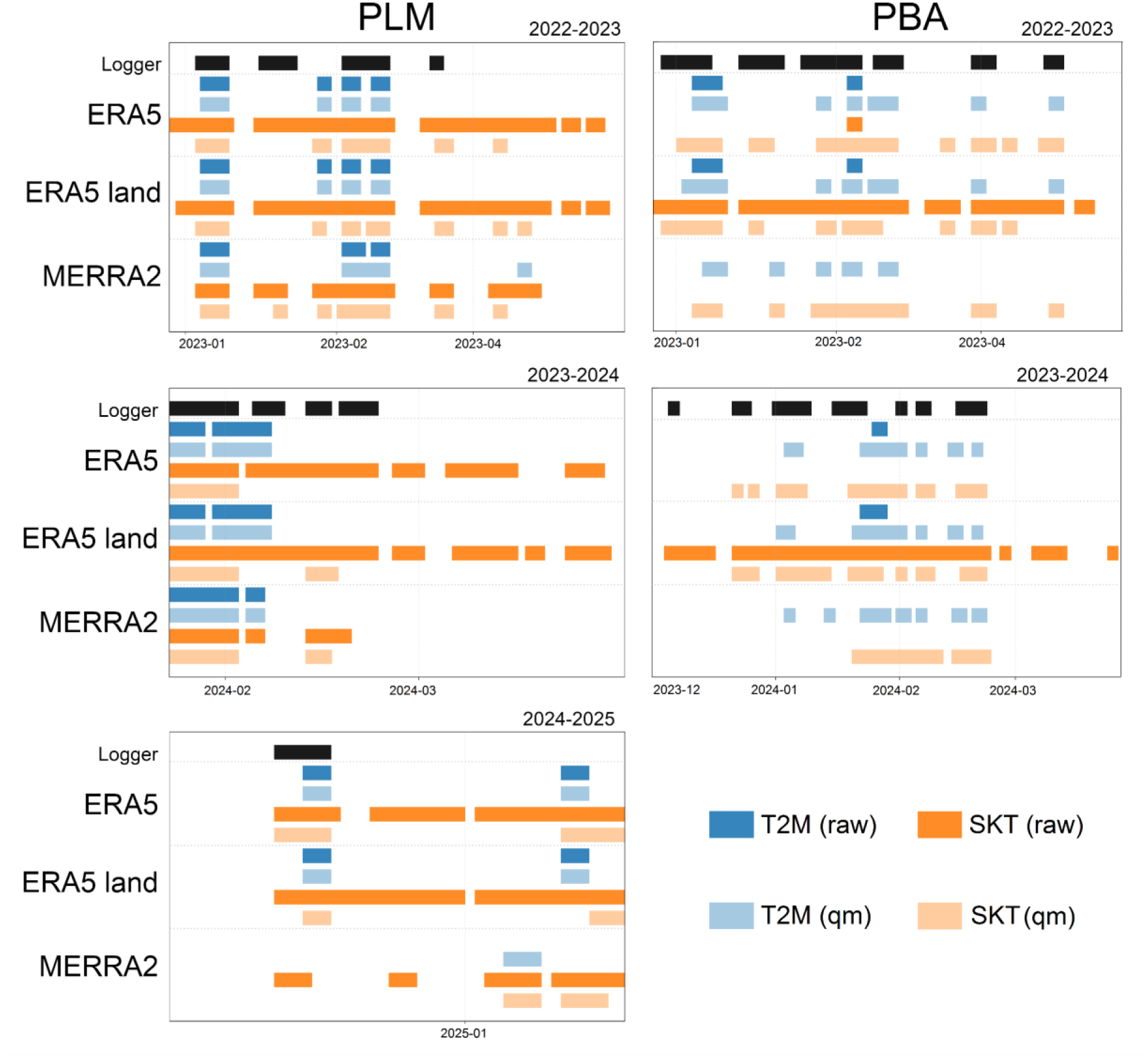
Timeline of heatwaves detected at PLM (left) and PBA (right) during the analyzed summer seasons. Each horizontal bar represents the duration of a heatwave event (temperature threshold ≥28.0 °C, duration ≥3 days) identified by the *in situ* logger (black) and by different reanalysis products (colored bars). Events are disaggregated by atmospheric variables: t2m (blue) and skt (orange). Solid dark colors indicate raw data, while lighter shades denote Quantile Mapping–corrected data.

At PLM, the raw skt products (ERA5, ERA5-Land, and MERRA-2) detected all heatwaves recorded by the loggers (100% detection rate; Fig. 5A). However, this apparent agreement was also associated with systematic over-detection, as ERA5 and ERA5-Land skt generated seven additional heatwaves each, and MERRA-2 skt produced four false positives (Fig. 6). Furthermore, ERA5 and ERA5-Land skt exhibited a propensity to overestimate the duration and intensity of heatwaves (Fig. 5B, C). Conversely, the t2m products demonstrated a comparatively diminished performance, identifying a mere 44–56% of the observed events and yielding a lower number of false positives. Following the implementation of QM correction, the propensity for over-detection among the skt products was generally mitigated, although in certain instances, detection rates underwent a decline. For instance, ERA5 skt QM declined from 100% to 55.6%, while ERA5-Land skt QM decreased from 100% to 77.8%, reducing false positives from 7 to 4. In general, the application of QM correction resulted in a more conservative heatwave detection outcome. This was characterized by a reduction in false positives and an enhancement of the realism of the detected patterns, particularly with regard to ERA5-Land skt.

### Historical trends in heatwave frequency

The trend analysis of heatwave metrics derived from ERA5-Land skt at PLM for the period 1960–2024, based on the non-parametric Mann–Kendall test, showed that the mean temperature during heatwaves increased significantly, with a Kendall τ of 0.175 (p = 0.0265; Fig. 7), corresponding to a Sen’s slope of +0.12 °C per decade. In contrast, the remaining metrics did not display significant temporal trends over the study period, including the annual number of heatwaves (τ = –0.098, p = 0.2392), the mean duration of individual events (τ = 0.053, p = 0.5042), and the mean maximum temperature during heatwaves (τ = 0.107, p = 0.1757).

**Fig. 7.**
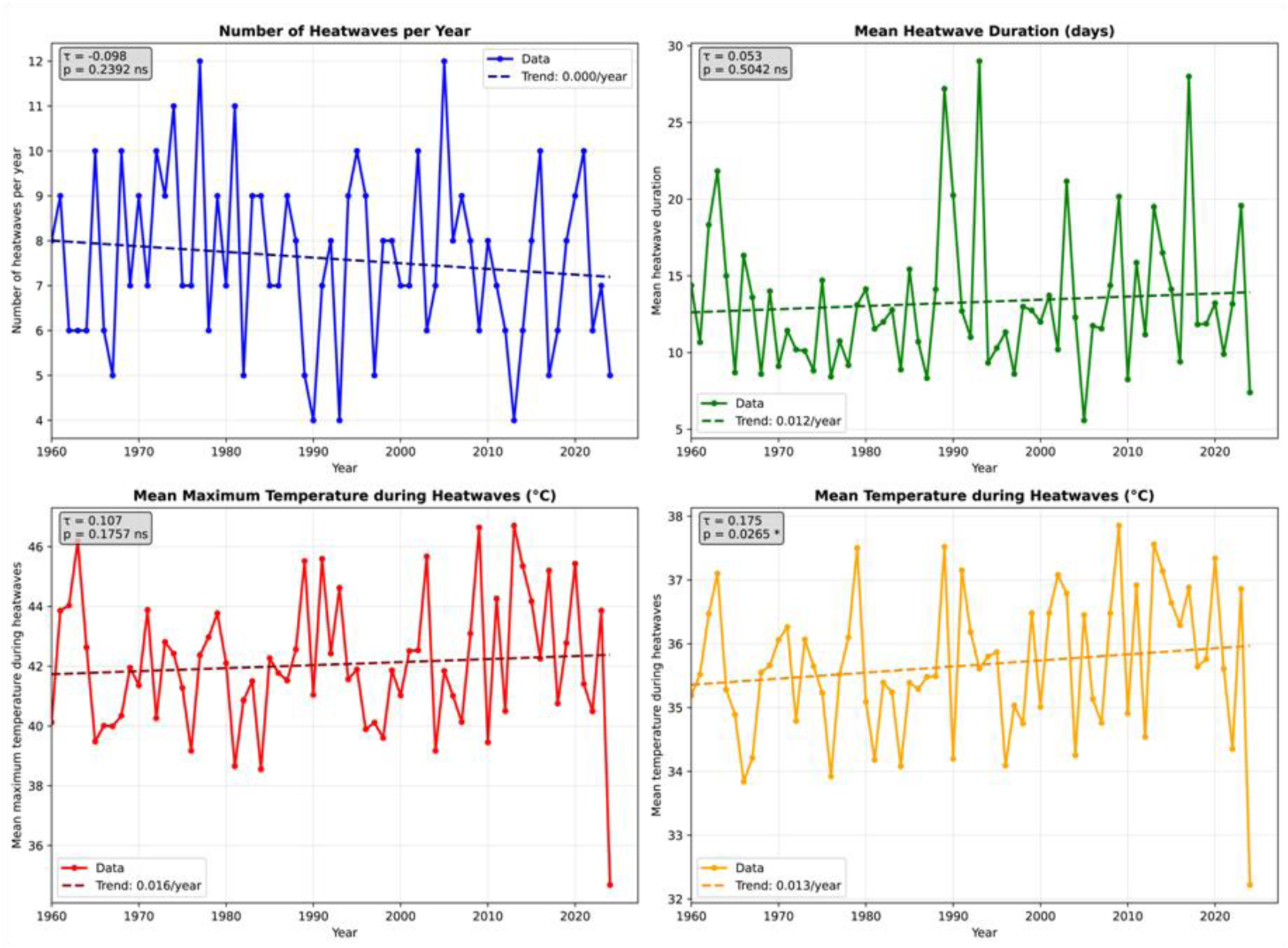
Trends in heatwave events detected at the PLM site from the ERA5-Land skt (1960–2024). Shown are: annual number of events (top left), mean duration (top right), mean maximum temperature during events (bottom left), and mean event temperature (bottom right). Dashed lines indicate the trend estimated using the Mann–Kendall test, with τ values and statistical significance reported in each panel

The comparison between the periods 1961–1990 and 1991–2020 revealed shifts in heatwave behaviour (Table 2). Although the annual frequency of events decreased by 7.8%, from 7.97 to 7.34 heatwaves per year, the mean duration of individual events increased by 10.4%, from 12.44 to 13.74 days. Both the mean maximum temperature during heatwaves and the mean event temperature increased by 1.2% (from 41.75 °C to 42.27 °C) and 1.1% (from 35.45 °C to 35.83 °C), respectively. Overall, the total annual number of heatwave days remained nearly unchanged, with only a slight variation of –0.2% between the two periods.

**Table 2.** Absolute and relative changes in heatwave characteristics between two standard climatological periods. Percentages indicate the relative change with respect to the 1961–1990 baseline.

| Metric | 1961–1990 | 1991–2020 | Absolute change | Relative change (%) |
| --- | --- | --- | --- | --- |
| Number of heatwaves per year | 7.77 | 7.50 | –0.27 | –3.4 |
| Mean heatwave duration (days) | 12.94 | 13.69 | +0.74 | +5.7 |
| Maximum temperature during heatwaves (°C) | 41.84 | 42.60 | +0.76 | +1.8 |
| Mean temperature during heatwaves (°C) | 35.43 | 36.03 | +0.60 | +1.7 |

The breakpoint analysis using the Pettitt test applied to the series of maximum temperatures (skt, ERA5-Land, 1950–2024) revealed a significant shift in the long-term behavior of the record (U = 4,748,774; p = 0.0025). The change point was detected on 24 September 2007, indicating an increase in the mean maximum temperature thereafter. Before the shift, the mean value was 24.56 °C; after the breakpoint, it rose to 25.29 °C, representing an increase of 0.73 °C in the mean maximum temperature. The Mann–Kendall test to the pre- and post-breakpoint intervals did not detect statistically significant monotonic trends either before the breakpoint (1950–2007: β = −0.014 °C yr⁻¹, p = 0.477) or after it (2007–2024: β = +0.060 °C yr⁻¹, p = 0.449).

## Discussion

In geographic areas where long-term *in situ* records of environmental parameters are deficient, validation and inter-comparison of reanalysis dataset can provide an adequate alternative for the detection of historical climatic variations on land. The intertidal zone, however, presents challenges as it is the interface between land and sea thus making it difficult to obtain accurate and reliable air temperature datasets. Our results show that reanalysis products can be a useful and trustworthy tool for recreating air temperature patterns on intertidal rocky shores where past *in situ* data is nonexistent. This was shown for SWA shores and has the potential to be applied in other data-poor coastal areas around the globe, albeit with previous ground proofing. More specifically, reanalysis performance depended on the chosen product, the type of series employed (skt vs. t2m), the application of a correction method and the site. In any case, correlations indicate robust relationships between reanalysis products and *in situ* logger measurements, providing confidence in the use of these datasets for regional climate and ecological studies. Products were also capable of detecting heatwaves, with little variation in their features for the period 1960–2024 at PLM. Thus, the validated reanalysis products proved to be a valuable resource for complementing local data and assessing long-term temperature patterns, providing key information to understand whether temperature variations in recent decades could have implications on ecological processes, for example on the changes in abundance of local species.

Using the global harmonic model TPXO9, thermal time series could be filtered to identify periods of air exposure in intertidal environments independently (Egbert & Erofeeva, 2002). This avoided the methodological circularity associated with approaches that infer emersion from abrupt temperature changes (Harley & Helmuth, 2003; Gilman et al., 2006). Within this framework, reanalysis products limited to emersion periods accurately replicated the variability of intertidal air temperature, demonstrating their usefulness for regional-scale ecological studies, particularly in coastal regions where *in situ* observations are scarce (Mislan & Wethey, 2011; Simmons et al., 2016). Hourly series of 2-m air temperature (t2m), representing near-surface atmospheric conditions, and skin temperature (skt), corresponding to the radiative temperature of the land surface, were extracted from the ERA5 reanalysis. These variables differ not only in their physical meaning but also in the processes they represent: while t2m integrates air temperature within a well-mixed layer above the surface, skt responds more directly to radiative heating and the thermal properties of the substrate. Consequently, the accuracy of the results depended strongly on the variable considered. As expected, skt showed better agreement with in situ logger records than t2m, since it more closely captures the rapid heating of exposed rocky surfaces during low tide, whereas t2m smooths temperature variability and tends to underestimate extreme values (Helmuth et al., 2002; Hersbach et al., 2020). Despite these robust relationships, absolute errors in raw reanalysis products remain high. This limits their application at fine spatial scales and highlights the need to complement these datasets with *in situ* measurements when assessing local ecological impacts (Judge et al., 2018; Helmuth et al., 2006).

Heatwaves have been associated with mass mortality events on intertidal shores worldwide (Garrabou et al., 2009; Hesketh and Harley, 2023; Thomsen et al., 2019). Some of these events included intertidal mussels, such as the scorched mussels *Perumytilus purpuratus* that dominate temperate Atlantic Patagonian rocky intertidal shores. As key foundation species, they form dense beds that support over 40 associated species (Silliman et al., 2011; Mendez et al. 2019; Mendez and Schwindt, 2024). Massive mussel losses, concentrated within Nuevo gulf, were registered after the 2019 austral summer (Mendez et al. 2021). Mussel cover remained high in nearby areas, including San José gulf. Mortality events were first related to increases in temperature and the higher frequency of occurrence of heatwaves, as has been observed for other intertidal mussels’ species (Harley, 2008; Seuront et al., 2019; Scrosatti 2024; White et al. 2021). For the first time in the area, in this work we obtained continuous *in situ* temperature data to which mussels are exposed to during low tide. Loggers registered heatwaves at the two sites, with a greater number of events recorded at PBA, in San José gulf. Detection rates of heatwaves with reanalysis products were high (more than 75 %) for both sites, with differences in performance again related with the product chosen and the use of correction methods. Of the evaluated indicators in the historical trend analysis at PLM, mean temperature during heatwaves was the only one that increased, whilst the annual number of heatwaves, the mean duration of events, and the mean maximum temperature remained unchanged during the 1960–2024 period. Furthermore, a change point was detected in September 2007, indicating an increase in the mean maximum temperature thereafter. Together, these results suggest that a warming climate alone wouldn’t be explaining *P. purpuratus* population losses in Nuevo gulf but rather it would be affecting mussels in combination with other local drivers (e.g. fishing, tourism, maritime traffic and other industrial activities). Nevertheless, aquarium experiments are currently being conducted to assess the lethal and sub-lethal effects that heatwaves can have on the species.

Looking in more detail at heatwave analyses, we observed that products performed differently between the sites analyzed. At PBA, raw products showed limited ability to identify heatwaves, underestimating their intensity frequency. In contrast, at PLM, skt variables allowed events to be detected adequately even without statistical correction. These differences between sites can be attributed to contrasts in coastal exposure and local conditions: PBA corresponds to an open point, more exposed to waves and wind, while PLM is located within a relatively closed gulf, with more muted thermal dynamics (Mislan & Wethey, 2011; Cavalleri et al., 2024). In this context, statistical correction using quantile mapping proved to be particularly relevant, as it reduced systematic biases and improved the representation of thermal extremes, as has been documented in other studies that applied this approach to climate variables in coastal environments (Enayati et al., 2021). In PBA, in particular, the correction was essential for improving the detection of heatwaves and bringing the reanalyses closer to the thermal conditions recorded *in situ*, while in PLM its impact was more limited, reflecting the initial better performance of the raw products. It is important to note, however, that the quantile mapping calibration was constrained by the short observational period available. This implies that the correction primarily improves consistency within the observed window rather than representing long-term climatological distributions. Future applications should therefore recalibrate the method using longer observational series, allowing a more robust characterization of the full range of thermal variability and extreme-event behavior.

The gradual rise in temperatures and the increase in the intensity and frequency of extreme events create the need to know in order to understand the climatic variations that have occurred in recent decades. This will allow for an accurate understanding of the biological and ecological consequences of regional thermal stress on organisms. This understanding is therefore determined by the availability of long-term thermal time series. Reanalysis datasets have arrived to fill this gap for geographic regions or particular environments, such as intertidal shores, where *in situ* observations are lacking. Our results allow us to infer information on temperature trends and heatwave features, helping elucidate the climate of the past in an ecologically relevant area of Atlantic Patagonia. Furthermore, we provide novel information on the most suitable reanalysis datasets for each purpose. The use of the information obtained after validation provides the broader community with tools to expand the understanding of other ecological phenomena that go well beyond the mussel mass mortalities included here. We encourage others to use this same dataset, or others derived from them, to answer their own questions about their species/environments of interest. With this alternative source of long data sets, past responses to changes in temperature can be now assessed. We recognize a low number of loggers deployed and the limited temporal coverage, however we believe that the information obtained is robust and novel and provides a promising starting point for climatological studies on intertidal environments.

## Acknowledgments

This research was partially funded by FONCyT PICT 2021 00114 to MMM.

